# Phoretic mites as benign passengers: no influence on mate choice in the burying beetle *Nicrophorus nepalensis*

**DOI:** 10.1101/2024.12.14.627428

**Authors:** Brendan Lan, Tanzil Gaffar Malik, Mu-Tzu Tsai, Yi-Ta Wu, Syuan-Jyun Sun

## Abstract

Mate choice is a fundamental aspect of sexual selection where the ‘chooser’ chooses a ‘courter’ by assessing a variety of traits that communicate potential fitness. However, the influence of interspecific interactions, such as symbiosis, on mate choice remains underexplored. We addressed this shortcoming with experiments on burying beetle *Nicrophorus nepalensis* and their interactions with phoretic mites *Poecilochirus carabi*. The mites can act either as mutualists or parasites depending on the presence of competitors and mite densities, thus potentially influencing mate choice. In a laboratory experiment, we presented female *N. nepalensis* with a range of natural mite densities: 0, 5, 10, or 20, with males carrying either 0 or 10 mites in an olfactory-based mate choice assay. Subsequently we allowed females to breed with their chosen male and all their mites before evaluating the fitness effects of the varying mite densities. We found that females across all mite densities had no preference for males with or without mites. In line with this, the mite densities had no effect on the brood size or the averaged larva mass. However, the mite densities per breeding cohort did positively affect the number of mite offspring. Our results suggest that mites act as benign passengers, not directly affecting mate choice or fitness.

## Introduction

Mate choice is a significant aspect of sexual selection that can influence the fitness of animals across sexually reproducing taxa (Andersson 1994; Rosenthal 2017; Ryan et al. 2007). Choosers assess potential mates’ fitness through a variety of traits that are indications of a potential mate’s fitness. Choosing an ‘incorrect’ mate can lead to a decrease in the potential fitness of the chooser. Most studies have focused on how mate choice is affected by traits that are either evaluated by the chooser or exhibited by the courter. For example, mate choice has been extensively studied in the context of ornamentation (Qvarnström et al. 2006), body size (López-Cortegano et al. 2020), age (Beck & Powell 2000), mating history (Aich et al. 2020; Arnqvist & Nilsson 2000), ability to win bouts (Emberts et al. 2018; Emlen 2008), and motor performances (Byers et al. 2010) across taxa.

However, many traits that affect fitness have been overlooked as potential influencers of mate choice. For instance, symbiotic relationships such as mutualisms and parasitisms can increase or decrease an individual’s fitness and have comparatively been seldom studied in the context of sexual selection. Among the studies that have examined how these symbiotic relationships may affect mate choice, parasitisms have been the obvious focus especially in the context of parasite avoidance (Beltran-Bech & Richard 2014; Cantini et al. 2024; Ehman & Scott 2002; Reyes-Ramírez et al. 2020) due to the more commonly intertwined relationship between host and parasite. Mutualisms, although they may confer significant fitness benefits, often do not directly affect the mate choice of either species involved as the associations are usually less intertwined than parasitisms. However, burying beetles and their phoretic mites that contextually act as protective mutualists or parasitic competitors (Beninger 1993; De Gasperin & Kilner 2015; Nehring et al. 2017; Sun & Kilner 2020; Wilson 1983; Wilson & Knollenberg 1987) present a system in which a complex symbiotic relationship has the potential to directly affect mate choice.

Burying beetles in the genus *Nicrophorus* are an intriguing subject of sexual selection studies as parents provide extensive care to their offspring - with many species showing high rates of offspring mortality in the absence of parental care, especially in early instar larvae (Trumbo 1992). In addition, individual *Nicrophorus* parents often work in tandem to compete against other carrion-breeders such as blowflies (Diptera: Calliphoridae). Together, this means that a potential mate is extremely important for the survival of one’s offspring and opens the door for sexual selection to be key in reproductive success. Indeed, mate choice studies in this genus examined the role that size (Beeler et al. 2002; Suzuki 2009), tendency to mate frequently (Hopwood et al. 2016), and rival presence (Suzuki 2009) play in *Nicrophorus* mate choice. An overlooked aspect of their mate choice that remains unexplored is whether the density of their phoretic mites affects their mate choice.

*Poecilochirus carabi* are common phoretic passengers on various *Nicrophorus* species, breeding in tandem with their host beetles on small carrion (Pukowski 1933). These mites have been experimentally shown to exhibit a mutualism-parasitism continuum in association with their burying beetle hosts (Sun & Kilner 2020). The burying beetles play the role of transportation, conferring a fitness benefit to *P. carabi* by acting as a flying shuttle to bring otherwise strictly cursorial mite passengers to the carrion on which both species breed. Once there, the mites will either aid beetles in competing with other carrion-breeders, thus increasing beetle fitness, or compete with these beetles and their larvae directly for carrion, thus lowering beetle fitness. However, in the absence of competitors, mites lower burying beetle reproductive success, particularly at high mite densities (Nehring et al. 2017; Sun & Kilner 2020). In addition, the presence of mites can help smaller beetles overcome their competitive disadvantage when competing with larger conspecifics for access to carrion, increasing their fitness (Sun et al. 2019). Consequently, an individual beetle’s phoretic mite density has direct fitness consequences for their offspring and thus, has the potential to act as a trait that affects mate choice.

In this study, we consider how the presence of naturally-occuring densities of *P. carabi* (hereafter ‘mites’) might influence mate preferences in female *N. nepalensis* using olfactory-based mate choice assays. Following mate-choice assays, female burying beetles were allowed to breed with their chosen males in order to assess the fitness consequences of their choice. Larvae number and larvae weight were measured as indicators of beetle fitness. The number of mites was also measured to determine the degree of mite success. We hypothesize that female burying beetles prefer male beetles without mites rather than males with mites, and that females have a stronger preference to males without mites when the females carry higher initial mite densities themselves. In addition, in line with their mate choices, resulting pairs’ total mite densities will have a negative relationship with burying beetle fitness.

## Methods

### Study species and field sampling

The experiments were conducted from April to May 2024. Both *N. nepalensis* burying beetles and *P. carabi* mites were cultured in the lab with ancestor individuals collected across northern Taiwan for genetic diversity. Maintenance of the beetle colonies is described elsewhere (*in revision*). Beetles were tested approximately two weeks following eclosion when they become sexually mature. From October 2023 to May 2024, we surveyed for the occurrence and natural densities of burying beetles alongside their mites using hanging traps, each baited with a commercial mouse carcass. In total, 1438 trapping events were conducted in northern Taiwan. Each fresh mouse carcass was left to decompose for four days prior to beetle and mite collection. Once captured, we separated the mites from the beetles using a fine brush and pairs of tweezers. We processed the beetles individually by sexing them and determining their body size by measuring the pronotum width to the nearest 0.01 mm with a vernier caliper. In addition, we counted all *P. carabi* mites associated with each beetle to determine mite density per beetle.

### Experimental setup

To set up the mate choice assay and the subsequent fitness assessment of breeding pairs, beetles were first measured upon sclerotization post-eclosion. Measurements of the pronotum width were used as a proxy for beetle body size as is the standard procedure (Hopwood et al. 2016; Jarrett et al. 2017). A total of 80 female beetles and 160 male beetles were used. Male beetles were matched into pairs that were from the same family and size-matched so that there was less than a 0.6 mm (0.18 ± 0.15 mm; mean ± SD) pronotum width difference between individuals. These matchings were done to minimize variations caused by genetics and by size. Following this, each of the resulting male pairs were matched with individual females, ensuring that the male pairs were not genetically related to the females. These trios were then randomly assigned to four treatment groups (n = 20 trios per treatment; Table 1). The densities of phoretic mites traveling on these beetles in situ varies, with previous sampling discovering that 84.4% of wild *N. vespilloides* carried between 0 and 20 P. carabi mites (Sun et al. 2019) and a similar distribution is found for *N. nepalensis*. To test whether various naturally-occurring mite densities affected female mate choice, we used four treatments: 0 mites, 5 mites (low density), 10 mites (medium density), 20 mites (high density) (Table 1). Each male within a matched male pair was either treated with 0 mites or 10 mites. Beetles that were assigned to be treated with mites were treated ca. two hours prior to mate choice assays. All beetles were fed small pieces of ground pork every 3-4 days prior to the trials.

**Table 1.** A summary of the treatment groups used in the mate choice studies and subsequent fitness assessments.

| Treatment | Female Beetles | Male Beetles | Resulting Mite Numbers per Cohort |
| --- | --- | --- | --- |
| 1 (n=20) | 0 mites | 0 mites | 0 mites |
|  |  | 10 mites | 10 mites |
| 2 (n=20) | 5 mites | 0 mites | 5 mites |
|  |  | 10 mites | 15 mites |
| 3 (n=20) | 10 mites | 0 mites | 10 mites |
|  |  | 10 mites | 20 mites |
| 4 (n=20) | 20 mites | 0 mites | 20 mites |
|  |  | 10 mites | 30 mites |
| Total: n=80 |  |  |  |

In order to simulate breeding conditions and encourage natural mate choice, a 22-30 g mouse carcass was introduced alongside each male in the holding chamber of the mate choice arena (Fig S2) similar to the preference trials used in (Delclos et al. 2021). These frozen-thawed mouse carcasses were sourced from a commercial feeder mouse seller (Mice Plus, Changhua, Taiwan) intended for consumption by pet reptiles. Each carcass was weighed prior to use and size-matched within one gram (0.36 ± 0.26 g; mean ± SD) to create size-matched pairs of carcasses. These carcass pairs were then randomly assigned to beetle trios and left outside for 30 hours in order to begin decomposition prior to mate choice assays making them more attractive to breed upon (Kalinová et al. 2009).

### Mate choice assay

An olfactory-based mate choice assay was conducted, because burying beetle mate choice and reproduction is likely olfactorily mediated as male burying beetles release pheromones (‘calling’) to attract females (Beeler et al. 1999; Trumbo & Eggert 1994) and other key behaviors such as locating a carcass (Kalinová et al. 2009) and nestmate recognition (Steiger et al. 2008) are also olfactory-based. In addition, the mate choice of females was studied rather than males, because female *Nicrophorus* species contribute more towards the parental care of offspring (Hwang & Lin 2013; Scott & Traniello 1990) indicating that they are likely to be the more ‘choosier’ sex. This is supported by the fact that females seem pickier in other *Nicrophorus* species as well, sometimes refusing to copulate with males (Suzuki 2000).

For each trial the female was first placed in the acclimation chamber while the two males and their respective mouse carcasses were placed in their own holding chambers (Fig S2a). The female was held in the acclimation chamber for at least five minutes before commencing the trial. After acclimation, the female was allowed to freely roam the arena by removing the removable door. During this time if the female made physical contact with a choice wall (Fig S2a), the male beetle and carcass in the associated holding chamber was determined to be her mate choice. If the female did not make contact with a choice wall within 20 minutes post-acclimation, she was determined to have not made a choice and was not allowed to breed. Throughout the entire trial including acclimation, air movement was powered by an air sampling pump (SKC XR 224-PCXR8, SKC Inc., USA) and volatiles were allowed to pass through the arena as indicated by Fig S2a. The airflow was measured to be 19 mL/minute. Each trial was done in the dark under red lights (Fig S2b,c) to simulate a nighttime environment and conducted at approximately 21°C. Mate choice assays were conducted at 5:30 pm each day as that is a time at which beetles are active. The arena was made from laser-cut acrylic. Up to four trials were conducted at the same time (Fig S2c).

### Fitness assessment

Upon the female beetle making a choice in the mate choice assay, the female, her chosen male, all mite passengers on both beetles, and the carcass in the holding chamber were removed from the mate choice assay. These cohorts were then allowed to breed on the carcass in a cylindrical plastic container (14.2 cm diameter x 6.3 cm height) with a soil depth of approximately two centimeters at a mean ambient air temperature of 17.8°C. These containers were darkened with an opaque covering to encourage natural breeding events of burying beetles.

Approximately 90 hours after the burying beetles were allowed to breed, the number of eggs were counted as a measure of fitness. However, to count the egg number without disturbing the parents, only the number of eggs that were visible from the bottom of the cylindrical container were counted as a proxy for the total number of eggs each female laid.

Upon larval dispersal, the brood size, the weight of each brood, and the number of mites were measured. Larval dispersal usually occurred approximately 10-11 days after the burying beetles were allowed to breed, however, some broods developed slower. As such these measurements of the late-dispersal broods were only conducted once larvae had reached 3rd instar and were ready to disperse. The number and weight of larvae were counted by hand and weighed with a scale to the nearest 0.001 g (Shimadzu; Model: ATX224R). A benchtop counter was used to determine the number of mites phoretic upon both of the parent beetles. This number was used as a proxy for the total number of mite offspring (the number of mites post-breeding), as the majority of the mite offspring tends to disperse with adults compared to the offspring (De Gasperin & Kilner 2015).

### Statistical analysis

To analyze female burying beetle mate choice between males with and without mites, we performed binomial tests using the ‘binom.test’ function for each mite density treatment (0, 5, 10, and 20 mites) of female beetle to determine if the proportion of females choosing males with mites was significantly different from 0.5, which would indicate a lack of preference.

To analyze the effects of mite density treatment and mate choice on the fitness consequences of beetles, we used a generalized linear mixed model (GLMM) to analyze the brood size with a negative binomial distribution to account for data overdispersion. Female mite density treatment (0, 5, 10, and 20 mites) and the female mate choice (males with or without mites) were included as categorical variables, whereas female body size and carcass mass were included as covariates. In addition, we analyzed the brood size in another model, in which we considered the total number of mites (mites on both the male and the female) as a continuous variable, wherein female body size and carcass mass were also included as covariates. To analyze the average larva weight per brood (hereafter ‘averaged larval mass’), we used a GLMM with a gaussian distribution. Similar to analyzing brood size, we included female mite density treatment and the female mate choice as categorical variables, female body size and carcass mass as covariates, and additionally we included the brood size as a covariate since brood size has been known to negatively determine averaged larval mass in burying beetles (Schrader et al. 2015). In addition, we analyzed the averaged larval mass in another model, in which we considered the total number of mites (mites on both male and female beetles) as a continuous variable, wherein female body size, carcass mass, and the brood size were included as covariates.

To analyze the effects of mite density on the fitness consequences of mites, we used a GLMM to analyze the number of mite offspring with a negative binomial distribution. Total number of mites was included as a continuous variable, whereas female body size and carcass mass were included as covariates. To determine the relationship of fitness consequences between beetles and mites, we used GLMMs to analyze the brood size and the averaged larval mass with a negative binomial distribution and gaussian distribution, respectively. In both models, we included the number of mite offspring, female body size, and carcass mass as covariates, and additionally included the brood size when analyzing the averaged larval mass.

All statistics were conducted in R statistical software version 4.1.2 (R Development Core Team, 2011). We evaluated all model residuals for normal distribution and overdispersion using the *DHARMa* package (Hartig & Lohse 2021). GLMMs were analyzed using the ‘glmer’ function in the *lme4* package (Bates et al. 2012). We obtained *p* values for the main effects using the ‘Anova’ function in the *car* package (Fox et al. 2015). In all GLMMs, the family origin of female beetles were included as a random factor to account for this variation since multiple females from the same families were used in the experiment.

## Results

Of 1438 trapping events, 528 traps caught at least one beetle. In these traps, the number of beetles per trap ranged from 1 to 18, with an average of 2.8 beetles per trap. Our field trapping further showed that 72% of *N. nepalensis* individuals carried at least one individual *P. carabi*. The number of mites per individual ranged from 0 to 166, with an average of 8 mites. The mean and frequency distribution of mite numbers were similar between males and females (mean: t = −0.61, *p* = 0.540; distribution: D = 0.05, *p* = 0.240; Fig S1).

There was no significant preference in female beetles for male beetles with or without mites across all female mite densities tested (0 mite: *p* = 0.238; 5 mites: *p* = 0.815; 10 mites: *p* = 0.804; 20 mites: *p* = 0.332; Fig 1). In line with this lack of significant preference in mate choice, different initial mite numbers of just females and total mite numbers in each cohort both had no effect on the brood size (χ^2^ = 1.13, d.f. = 3, *p* = 0.770; Fig 2a; χ^2^ = 0.06, d.f. = 1, *p* = 0.807; Fig 2b) and the averaged larval mass (χ^2^ = 4.99, d.f. = 3, *p* = 0.173; Fig 2c; χ^2^ = 3.33, d.f. = 1, *p* = 0.068; Fig 2d). Although the starting number of total mites in each breeding cohort did have a positive relationship with the number of mite offspring (χ^2^ = 4.35, d.f. = 1, *p* = 0.037; Fig 3), the mite offspring number did not explain beetle larvae fitness as the number of individuals in a brood (χ^2^ = 1.71, d.f. = 1, p = 0.191; Fig 4a) or averaged larval mass (χ^2^ = 0.583, d.f. = 1, p = 0.445; Fig 4b).

**Fig. 1.**
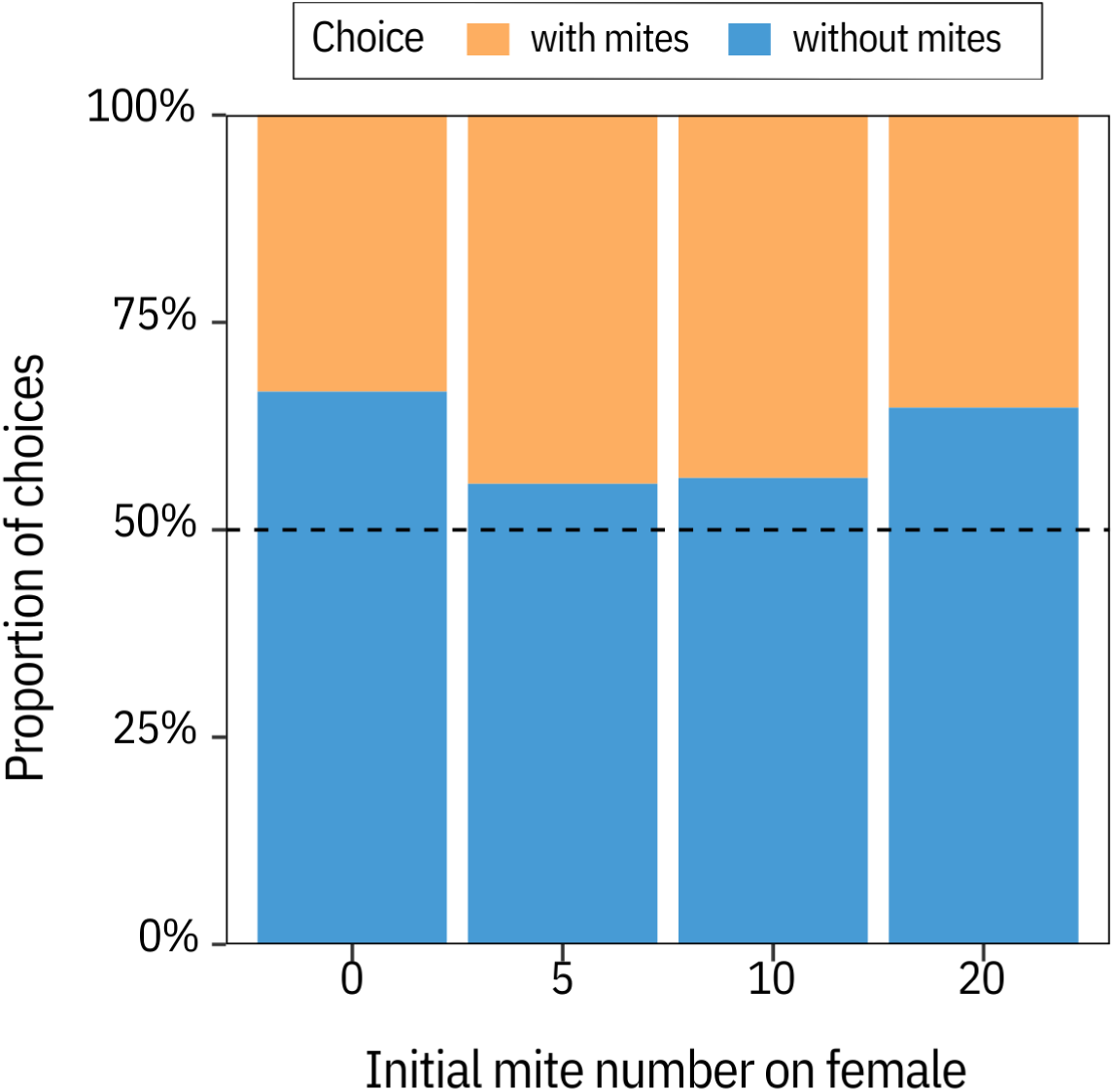
The number of mites on the female had no effect on her choice of male mate in all mite density treatments. The dashed line indicates no preference.

**Fig. 2.**
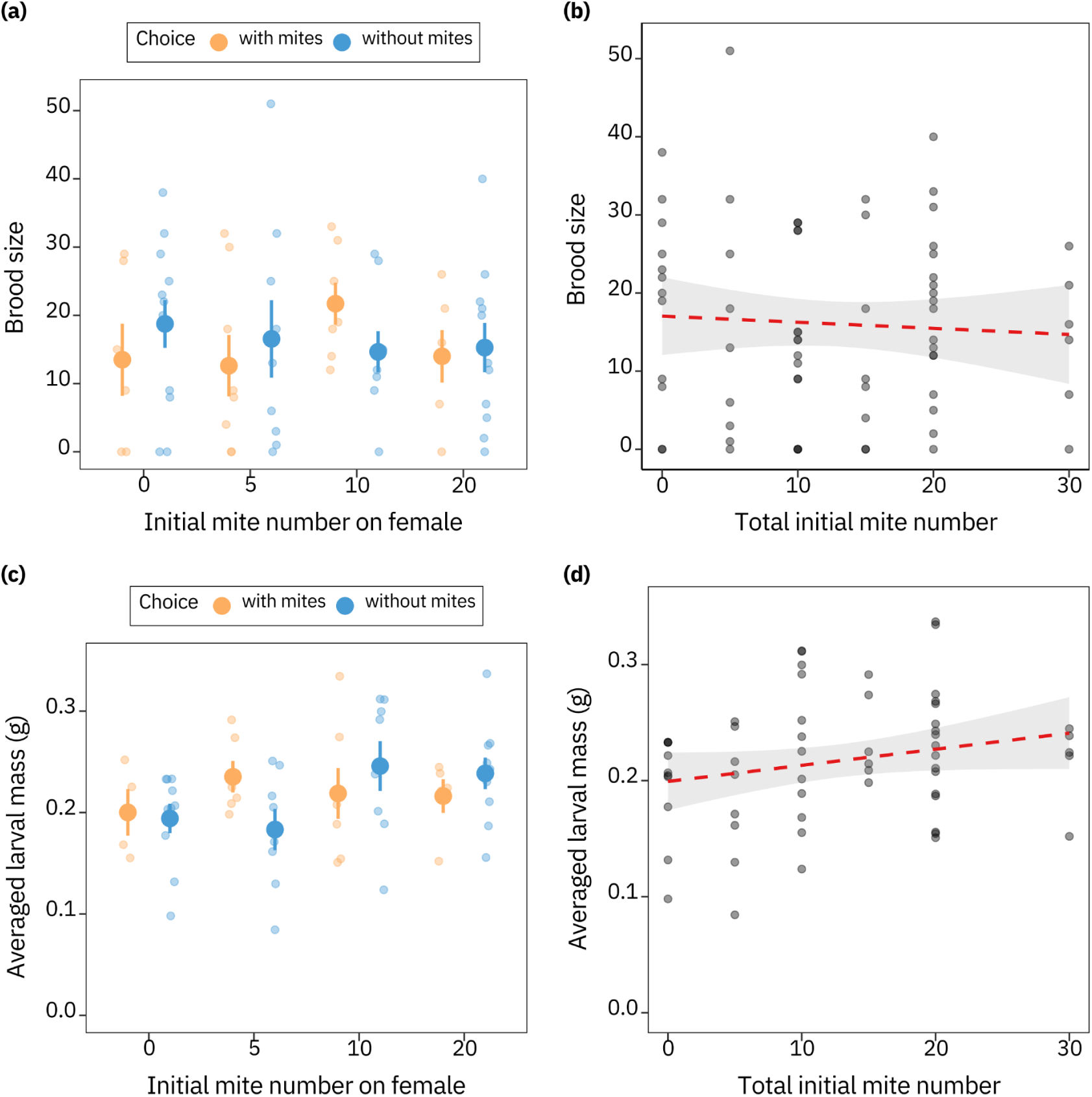
The effects of the initial mite number on just the female (a,c) and the total mite number of each breeding cohort (b,d) on the brood size (a,b) and averaged larval mass of each brood (c,d). a) The effect of the initial mite number of just the female beetle on brood size. (b) The effect of total mite number of each breeding cohort on brood size. (c) The effect of initial mite number of just the female beetle on the averaged larval mass of each brood. (d) The effect of total mite number of each breeding cohort on the averaged larval mass of each brood.

**Fig. 3.**
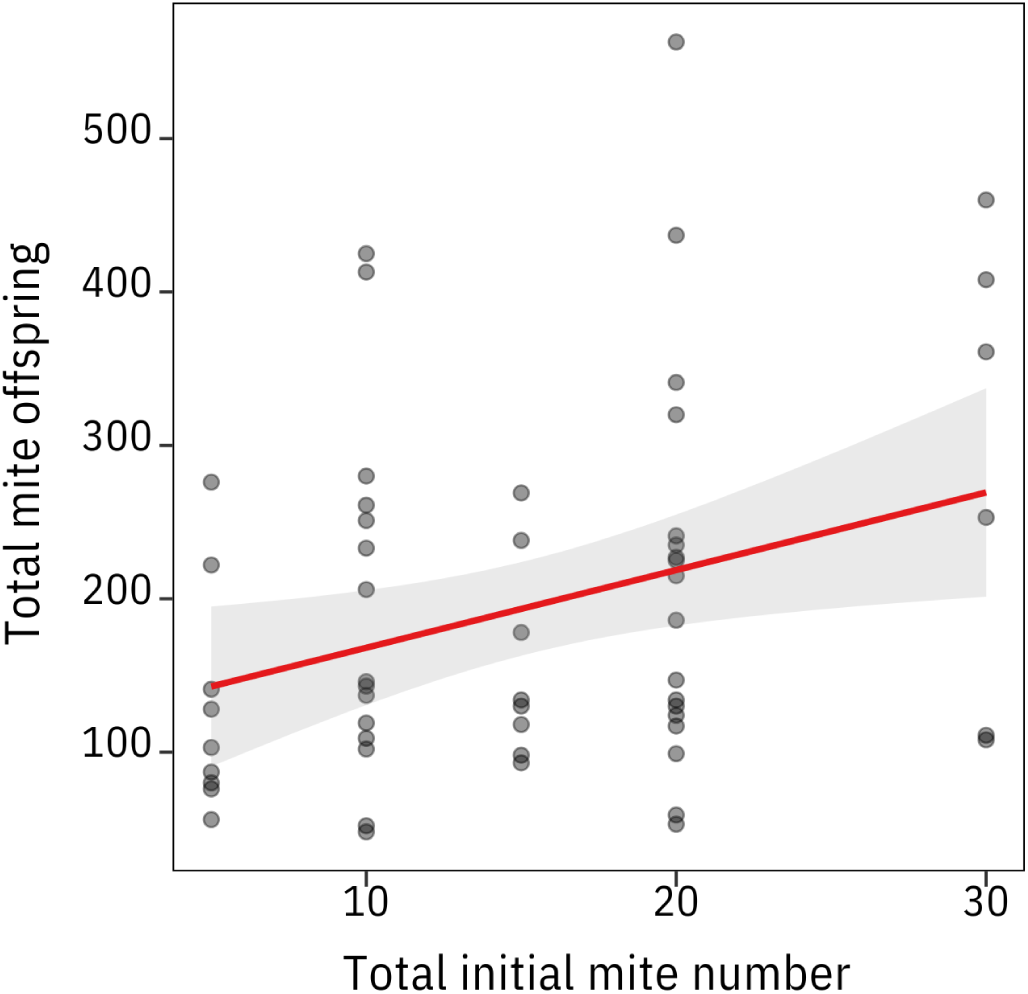
The total initial number of mites per breeding cohort had a positive effect on the total number of mite offspring. The solid line indicates statistically significant relationship from GLMMs, whereas the shaded area represents 95% confidence interval.

**Fig 4.**
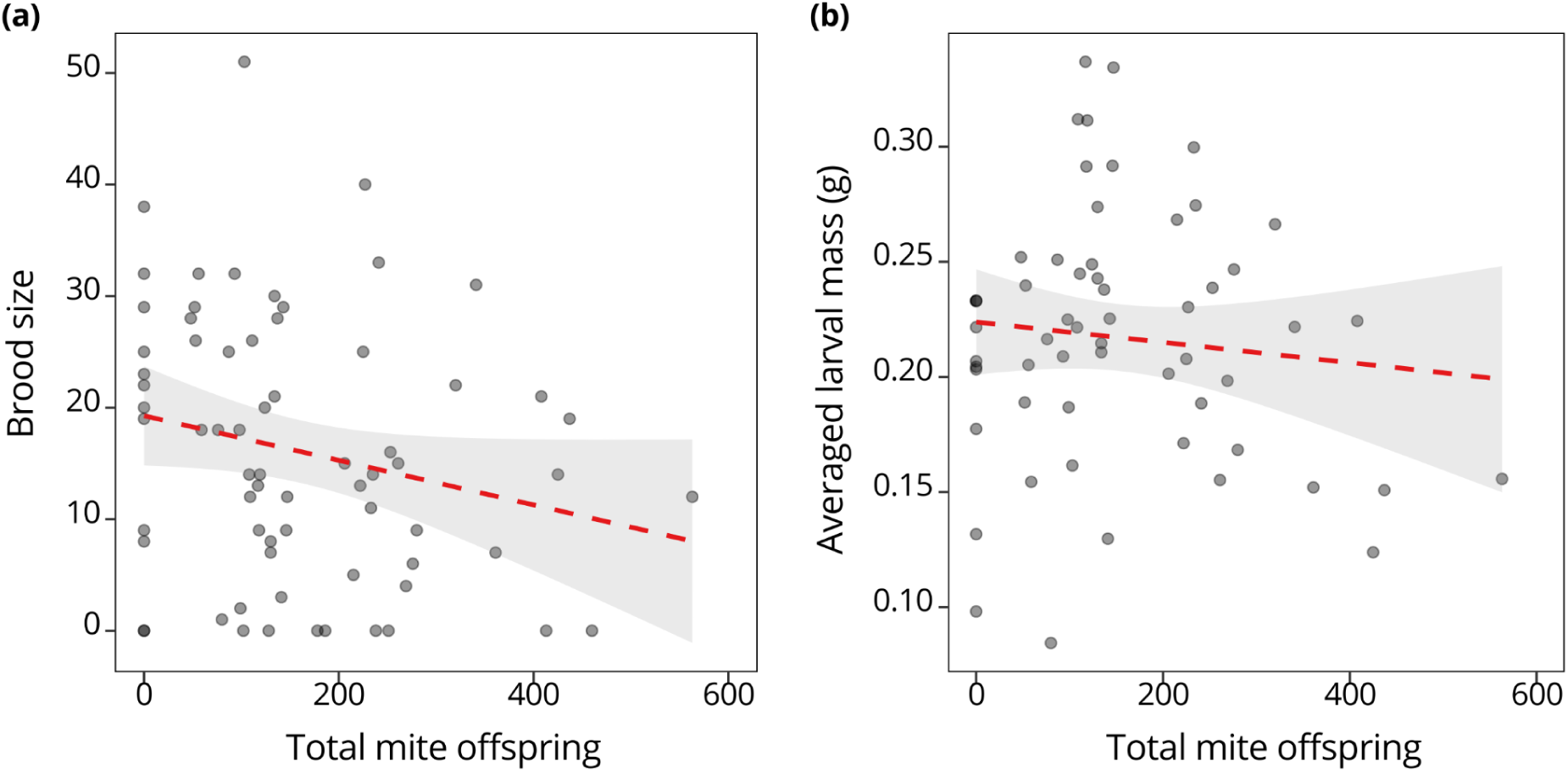
The total number of mite offspring resulting from each breeding cohort had no effect on brood size (a) or the averaged larval mass in each brood (b). The dashed lines indicate statistically non-significant relationships from GLMMs, whereas the shaded areas represent 95% confidence intervals.

## Discussion

Our findings suggest that mites in naturally-occuring densities do not affect burying beetle mate choice or fitness and that these mites are benign passengers in the absence of competitors. Although the starting mite densities of each pairing did positively affect the number of total mite offspring per cohort (Fig 3), burying beetle fitness was not affected (Fig 4). However, this relationship is likely modulated by other biotic and/or abiotic factors, which, in turn, could influence mate choice in burying beetles. Based on these results, we conclude that mites are mostly benign passengers that have little influence on mate choice in *N. nepalensis*.

Mate choice in *Nicrophorus* has been investigated broadly in the past as parent beetles of many species in this genus provide extensive biparental care. Thus, the choice of a potential mate is extremely important. As mentioned above (see ‘Introduction’), mate choice studies in this genus have often focused on the role that size plays in burying beetle mate choice (Beeler et al. 2002; Suzuki 2009). Beetle size is an indicator of fitness because larger beetles win fights for access to carrion more often (Otronen 1988), and because larger beetles are more fecund (Scott 1998), thus size has the potential to affect fitness. The presence of mites can also affect fitness both positively and negatively. Mites contextually increase beetle fitness by both aiding smaller beetles in intraspecific fights (Sun et al. 2019) or outcompeting blowflies on carrion (Sun & Kilner 2020). Mites can also decrease beetle fitness by competing directly with beetle larvae in the absence of competitors (Nehring et al. 2017; Sun & Kilner 2020). This investigation is the first which investigates whether these phoretic mites with the potential to modulate beetle fitness affects mate choice.

Previous work found that mites acted as parasites in the absence of other competitors (Nehring et al. 2017); however, in this investigation, mites failed to impact burying beetle fitness. This discrepancy is likely due to one or both of the following factors: the temperature our cohorts were reared in and the species used for this experiment.

When considering temperature, this investigation allowed cohorts of burying beetles and mites to breed in incubators that were kept at a mean of 17.8°C. In Sun & Kilner (2020), three temperatures were used to rear beetles with varying numbers of mites: 11°C, 15°C, and 19°C. The beetles in the 19°C rearing treatment seemed to show the least negative fitness effect as mite density increased. Nehring et al. (2017) reported a negative relationship between mite number and burying beetle fitness at 20°C. Additionally, both studies used mite densities well within the range of the current investigation: 0, 10, and 20 mites per beetle pair in Sun & Kilner (2020) and 10 mites per pair in Nehring et al. (2017). Our current investigation had cohorts with 0, 5, 10, 15, 20, and 30 mites (Table 1). Thus it is unlikely that mite densities, at least within most naturally occurring contexts, modulate this relationship. Taken together, it is probable that this association between burying beetles and mites, with respect to reproductive success, is quite sensitive to temperature, ranging from a commensal association to a parasitic one with mites negatively affecting burying beetle fitness at the two ends of the temperature gradient.

In addition, to our best knowledge, the current study is the first empirical investigation into the mate choice and mite associations of *N. nepalensis*. It is possible that mites affect *N. nepalensis* differently than other *Nicrophorus* species, especially given differences shown in this relationship with the sympatric *N. vespilloides* and *N. vespillo* (Nehring et al. 2017) as well as between some *N. tomentosus* and other American *Nicrophorus* species (Wilson & Knollenberg 1987). More experimentation could be done to test the effect of mites on burying beetle fitness across a larger temperature range and across various *Nicrophorus* species, especially those of varying body sizes. If mites do affect mate choice in *Nicrophorus* in parasitic contexts, they likely complicate mate choice as previous work investigating the effects of parasites on mate choice have had quite varied results across systems (Cantarero et al. 2022; Ehman & Scott 2002; Reyes-Ramírez et al. 2020).

Although mites seem to be protective mutualists of burying beetles, this relationship only occurs in the presence of competitors such as blowflies (Sun & Kilner 2020; Wilson 1983; Springett 1968). It is possible that, in the presence of blowflies, burying beetles will preferentially choose mates with more mites to increase their fitness, because Sun & Kilner (2020) showed that as mite numbers increased, burying beetle fitness also increased in the presence of blowfly competitors. This relationship was reported to be affected by temperature (Sun & Kilner 2020), and thus temperature seems to be key to this symbiosis, both in the presence and absence of competitors.

From the perspective of the mites, it is interesting that an increase in the initial mite numbers of each cohort had a positive effect on their reproductive success (Fig 3), especially given that an average of 8 mites were found on wild beetles (Fig S1) and that we used initial mite densities up to 30 in each cohort (Table 1). This indicates that the range of initial mite densities tested do not reach the carrying capacity of the carcass in our conditions, even with burying beetles breeding upon them in tandem. This is likely because wild mites experience a wider range of initial mite numbers than tested in this investigation given that more than two beetles are often attracted to and breed upon one carcass as indicated by our field trapping results. Further increasing the initial number of mites would likely negatively affect mite reproductive success and potentially negatively affect burying beetle fitness.

Overall, this investigation presents evidence that naturally occurring densities of *P. carabi* are benign, commensal passengers to *N. nepalensis* with respect to reproductive success in the absence of other competitors (Fig 2) and that these mites do not affect *N. nepalensis* mate choice (Fig 1). Even with the positive effect of starting mite densities increasing mite offspring (Fig 3), burying beetle fitness remained unaffected (Fig 4). These fitness effects are likely modulated by temperature and the presence of competitors, as is the case with other *Nicrophorus* species (Sun & Kilner 2020; Nehring et al. 2017). However, whether mate choice in the presence of mites is affected by a shift in their contexts, such as different temperatures and/or the presence of competitors, that cause mites to act as parasites or as mutualists remains to be seen.

## Supplementary Figures

**Fig S1.**
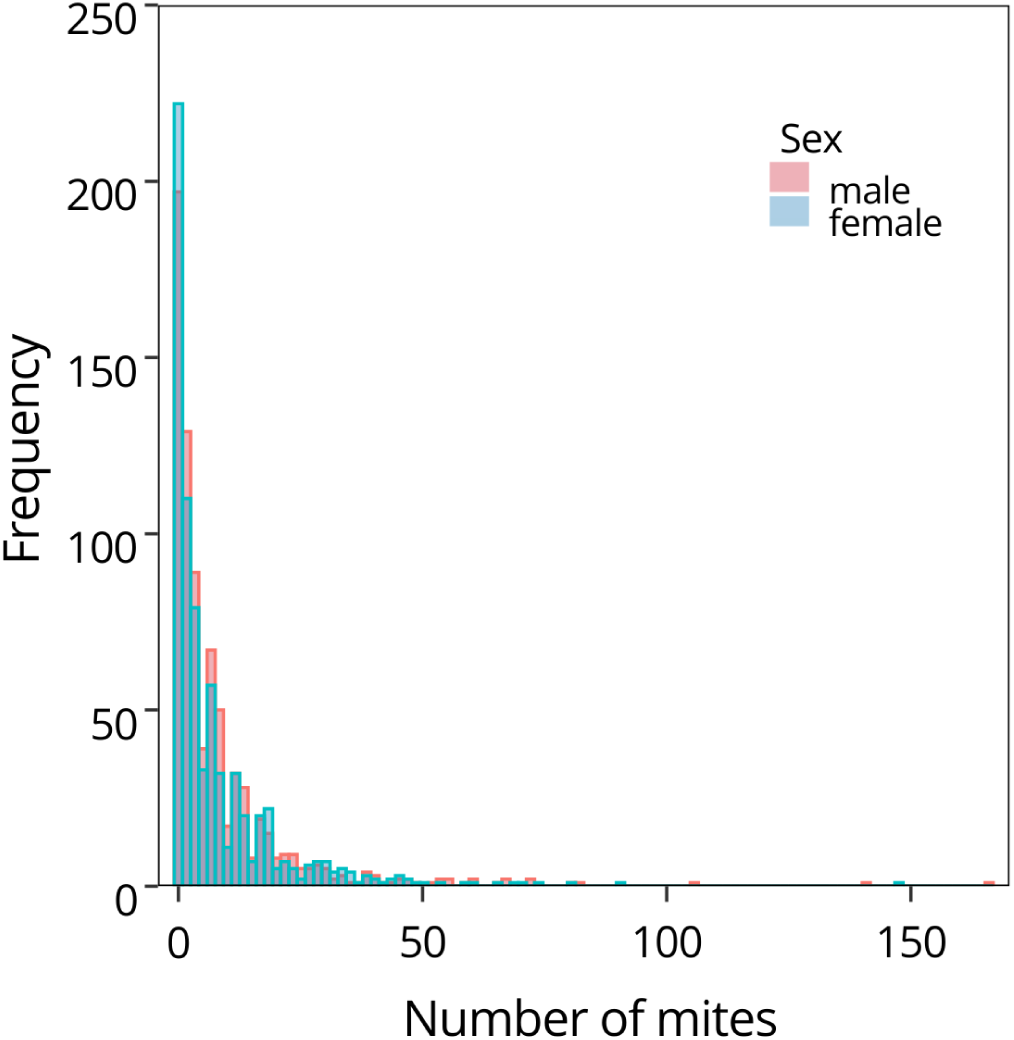
Histogram depicting results from counting the mites on field-trapped burying beetles. The mite number per individual ranged from 0 to 166, with an average of 8 mites.

**Fig S2.**
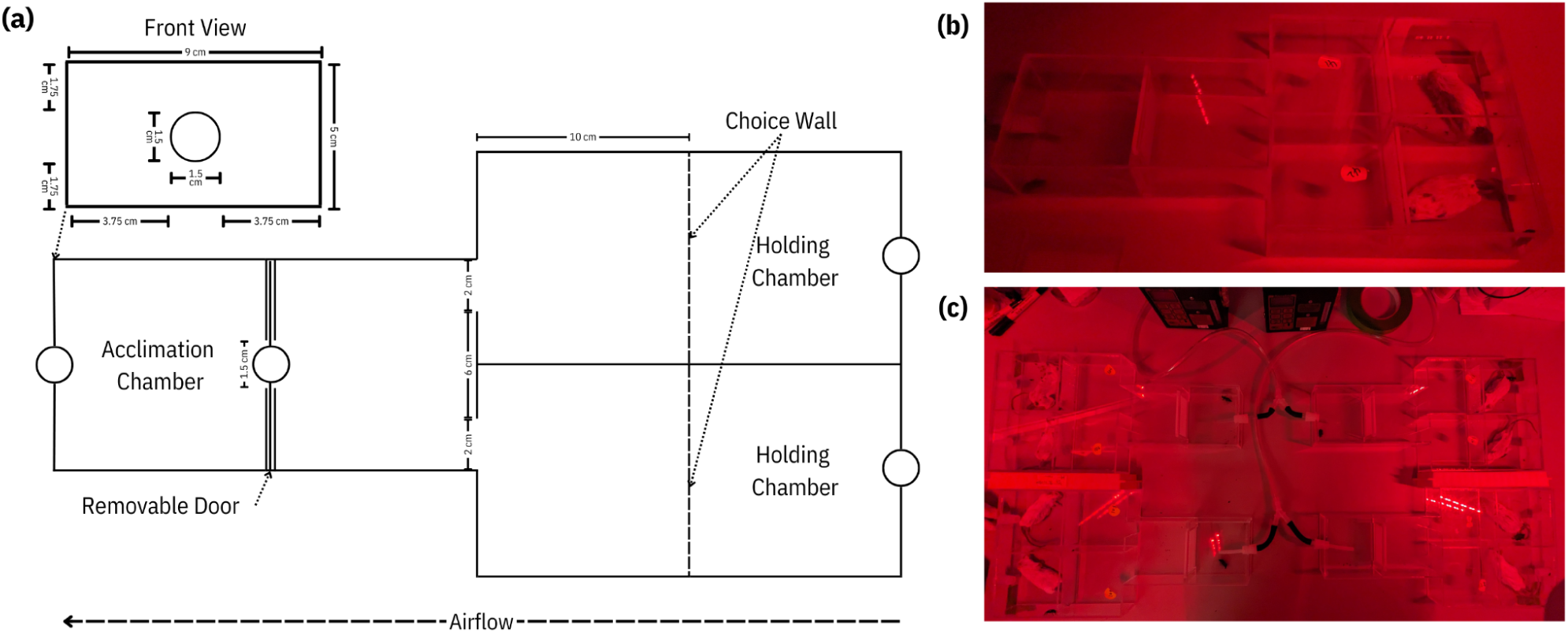
(a) Diagram of the mate choice arena depicting the Acclimation Chamber, Removable Door, Choice Walls, and Holding Chambers. The choice wall is perforated as indicated by the dashed line. Airflow moves as indicated by the dashed line arrow. The removable lid is not depicted here. (b) A photo of a mate choice arena in use with the female in the acclimation chamber, and both males with their respective carcasses in their holding chambers. (c) A photo of four mate choice arenas being utilized at once. In (b,c), the experiments were conducted under red light conditions.

